# Beauty-in-averageness and its contextual modulations: A Bayesian statistical account

**DOI:** 10.1101/360651

**Authors:** Chaitanya K. Ryali, Angela J. Yu

## Abstract

Understanding how humans perceive the likability of high-dimensional “objects” such as faces is an important problem in both cognitive science and AI/ML. Existing models of human preferences generally assume these preferences to be fixed. However, human assessment of facial attractiveness have been found to be highly context-dependent. Specifically, the classical Beauty-in-Averageness (BiA) effect, whereby a face blended from two original faces is judged to be more attractive than the originals, is significantly diminished or reversed when the original faces are recognizable, or when the morph is mixed-race/mixed gender and the attractiveness judgment is preceded by a race/gender categorization. This effect, dubbed Ugliness-in-Averageness (UiA), has previously been attributed to a disfluency account, which is both qualitative and clumsy in explaining BiA. We hypothesize, instead, that these contextual influences on face processing result from the dependence of attractiveness perception on an element of statistical typicality, and from an attentional mechanism that restricts face representation to a task-relevant subset of features, thus redefining typicality within that subspace. Furthermore, we propose a principled explanation of why statistically atypical objects are less likable: they incur greater encoding or processing cost associated with a greater prediction error, when the brain uses predictive coding to compare the actual stimulus properties with those expected from its associated categorical prototype. We use simulations to show our model provides a parsimonious, statistically grounded, and quantitative account of contextual dependence of attractiveness. We also validate our model using experimental data from a gender categorization task. Finally, we make model predictions for a proposed experiment that can disambiguate the previous disfluency account and our statistical typicality theory.

## 1 Introduction

Humans readily express liking and disliking for complex, high-dimensional “objects”, be they faces, movies, houses, technology, books, or life partners, even if they cannot verbalize exactly why. Understanding how these preferences arise is important for both cognitive science, and for AI systems that interact with humans. In particular, face processing presents a prime case study for complex information processing in humans. Human, including very young babies [17], readily perform sophisticated computational tasks based on a brief glimpse of a face, such as recognizing individuals, identifying emotional states, and assessing social traits such as attractiveness [9]. The last phenomenon has obvious impact on real-life decisions such as dating, employment, education, law enforcement, and criminal justice [38].

Existing models of human preferences, in both machine learning and cognitive science, have generally assumed social processing of faces (e.g. attractiveness judgment) to be a fixed function of the underlying face features [14, 34, 37, 39, 36]. However, a series of recent experiments have indicated that it is not a fixed process, but rather a fluid one depending on what other face-processing task the observer is also performing. Specifically, these experiments show that a classical phenomenon known as beauty-in-averageness (BiA), where blends of multiple faces are reliably found to be more attractive than the originals [16], can be diminished or even reversed (termed Ugliness-in-Averageness or UiA), when the facial blends are created from recognizable faces [11], or when attractiveness judgment of a mixed-race/mixed-gender morph is preceded by a racial/gender categorization task [12, 28].

The dominant account for the UiA findings is the disfluency account, which posits that the difficulty of categorizing (e.g. a biracial face) reduces perceived attractiveness by inducing a negative affect that is generalized to overall liking of the object [11, 12]. However, this account is not only qualitative, but also fails to satisfactorily explain the original finding of BiA when there is no categorization task. Instead, BiA has been explained by quite separate accounts, such as arising from an evolved preference to identify healthy mates [16, 10].

In this work, we propose a parsimonious, statistically grounded account of contextual dependence of human attractiveness judgment. Specifically, we propose that facial attractiveness depends on an element of statistical typicality, and that UiA effects arise when attentional mechanisms restrict face representation to a task-relevant subset (subspace) of features, thus redefining statistical typicality within that subspace. Furthermore, we propose the brain by default maps the input stimulus to a categorical prototype [24] and larger prediction errors [30], between expected featural properties of the categorical percept [24] and the actual sensory properties of the stimulus, induces a larger “cognitive cost” corresponding to greater encoding or processing cost. This increased cognitive cost, we propose, is what drives the lack of attraction of statistically atypical objects.

The rest of the paper is organized as follows. In section 2, we formally define statistical typicality and discuss the links to predictive error and affect. In section 3, we model the effects of attention, followed by proof-of-concept simulations in section 4 using an expository abstract model in 4.1 as well as a data drive-model of face space 4.2. In section 5, we validate our model using data from a gender categorization experiment. We propose a test to disambiguate the differences between the disfluency and statistical typicality based account in section 6 followed by discussion on limitations of our model and future work in section 7.

## 2 Statistical Typicality, Predictive Error and Affect

We assume an internal *d*-dimensional psychological face space *Χ* [40, 34, 27], in which any face *x* = (*x*_1_,…, *x_d_*) can be represented. We also assume that this face space is endowed with a statistical distribution *p_Χ_* (*x*) learned from the environment [6]. We will call *p_Χ_* (*x*), which may generally be a complex mixture distribution, as the *recognition model* and is the person’s *assumed* generative model of faces, the statistics of which may not necessarily correspond to generative distribution of faces in their environment due to mis-specified beliefs about the environment, incomplete information, varying importance of different faces or computational constraints.

Typicality is an important concept in cognitive psychology [32, 24] and high typicality of objects occurring either due to natural variance in features or induced by digital manipulation (e.g. by creating composite blends or by distorting objects toward the population prototype) has been shown to be associated with positive affect [16, 10, 31]; this is the classic BiA. Further, adding more objects to the composite generally makes the effect stronger. These findings motivate our definition of the statistical typicality of a face *x*, which we operationalize simply as log *p_Χ_* (*x*|*c*), the log-likelihood of *x* under the recognition model conditioned on task context *c*. Assuming that visual perception of faces is an inference process [42, 21, 15] employing the recognition model, the statistically optimal prediction *x*\* is simply the mean/prototype of the distribution *p_Χ_* (*x*|*c*) (under the loss function *d*(*x, x*\*) = (*x* − *x*\*)^2^). In the case of BiA, the context *c* is empty.

From an information theoretic perspective [2, 5], the more atypical a face *x*, the larger the cost (in bits) of encoding the prediction error/deviations from *x*\*. Although we will not work at the implementation level of abstraction in the parlance of Marr’s three levels of analysis [22] and instead focus on the computational level, these abstract ideas find parallels in the neurally plausible framework of predictive coding [30]. In the predictive coding framework, *x*\* corresponds to a top-down prediction, while the prediction error finds a ready parallel in feedforward residual errors. We propose that the increased cognitive cost required to encode the deviations of atypical faces from the prototype *x*\* induces a negative affect, providing a normative account of BiA. While there is an abundance of work suggesting that prototypes are processed with less cognitive effort and higher speed [29, 43, 35, 33], these have generally been in the context of categorization tasks and do not directly apply when there is no categorization task, such as in BiA.

## 3 Attentional Effects

In this section, we show that attentional effects modeled as task-relevant subspace projections can account for UiA in blends of known/recognizable faces as well as in race/gender categorization tasks using mixed-race/mixed-gender blends respectively. Perceptual tasks are often performed in a low-dimensional task-relevant subspace [7] focusing on informative dimensions in high dimensional sensory data using attentional modulation that is top-down goal-directed [26] or bottom-up saliency based [13]. We therefore model attentional modulation as projection to a task-relevant subspace 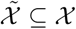. Denoting the projection of a face *x* into a subspace 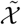 as *x̃*, we redefine statistical typicality in the subspace 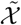 as 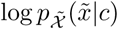, the log-likelihood of *x̃* under the recognition model constrained to subspace 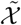, conditioned on task context *c*.

### Top-down attentional modulation

A categorization task such as gender or race categorization is modeled as a projection to a race/gender informative subspace 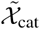. People often use category membership to predict features and reason about members of the category [20, 25]. Accordingly, statistical typicality in this case is 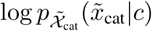, where the context *c* in this case is the a posteriori most probable category. Mixed-race/mixed-gender blends are statistically atypical of both categories inducing a negative affect.

### Bottom-up saliency based attentional modulation

It has been suggested that people encode familiar faces using features that are most distinctive/salient as this is not only computationally efficient but may also boost recognition [23]. Accordingly, we assume that familiar faces are represented using the *s* most salient features. More concretely, we assume *x_f_* is represented by its veridical value from the “true” generative model, if it is among the top *s z*-scored dimensions and 0 otherwise. We propose that a blend of recognizable/familiar faces *x* induces an implicit categorization in the subspace 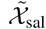 spanned by the distinctive features of the blended image via bottom-up saliency based attentional mechanisms. Similar to the context in explicit categorization tasks, *c* is the a posteriori (i.e., after classification) most probable identity, so that that statistical typicality in this case is 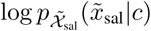 and is low for the blend *x* for all possible values of the a posteriori categories.

## 4 Simulations

We will first present a very simple, abstract model in section 4.1 that captures both BiA and as well as UiA in various contexts. The simplicity of this model is deliberate in that it is meant to be both expository as well as demonstrating the generality of our proposal, since BiA and UiA are not specific to faces [10, 43, 41]. In section 4.2, we use a data-driven face space representation for further validation.

### 4.1 Abstract Model

#### 4.1.1 Recognition Model

We assume that humans internally represent each face *x* = (*x*_1_,…, *x_d_*) ∈ ℝ*^d^* as generated from a mixture of Gaussians, whereby the components can either correspond to well-known faces *f_i_* (assume *K* of these) or demographic subgroups *h_r_* (assume *G* of these, e.g. gender, race),

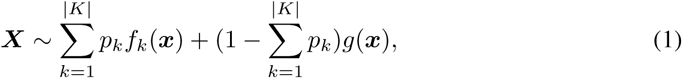

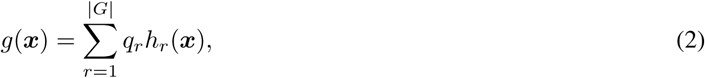

where *h_r_*(***x***) = *𝒩* (***x**; **μ**_r_*, **Σ**_*r*_), *f_k_*(***x***) = *𝒩* (***x***; ***μ**_k_*, **Σ**_*k*_) and 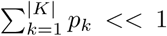 as the number of known faces should be much fewer than unknown faces. We assume that the statistics of the mixture components *h_r_* differ only in a small number of dimensions, 1,…, *d*_race_ and have the same statistics on the other *d*_other_:= *d* – *d*_race_ dimensions. Specifically, we assume 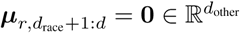 and

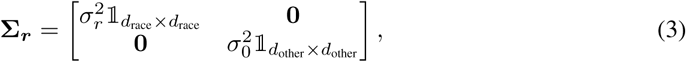

where 𝟙_*n*×*n*_ is an identity matrix of dimensions *n* × *n*. For simplicity, we assume |*G*| = 2 and set ***μ***_1_ = –***μ***_2_ = ***μ***, where 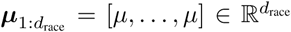. We also set the prior/mixture probability distribution *q* to be uniform.

##### Approximation

Note that since the statistics of *h_r_* differ only in a small number of dimensions *d*_race_ << *d*, the mixture 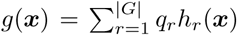 is well approximated by *g̃*(***x***) = *𝒩* (***x**; **μ***_0_, **Σ**_0_), where *μ*_0_ = 0 ∈ ℝ*^d^* and 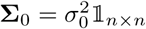 and can be assumed to be used to perform inference except when demographic features bear relevance, thus simplifying computations and representation.

##### Salient feature representation

The mixture components *f_k_* represent known/recognizable faces with variability arising naturally out of perceptual and representational noise as well as some inherent variability present in facial features. For each face *k*, we assume subjects encode/represent only *s* distinctive features (relative to the assumed generative distribution) as described in section 3, denoted by 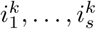 (the variance along these dimensions is denoted as 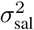) and assume the same statistics along other dimensions as *g̃*(***x***), the approximate, assumed generative distribution for a generic, unfamiliar face.

#### 4.1.2 Simulation Results

##### BiA

A general trend of increasing attractiveness with increasing number of images in a composite has been noted in [31]. This is naturally captured by increasing typicality of the composite with increasing number of constituent face images (Figure 1b). Additionally, the model also captures “more blended” composites of two images being more attractive than “less blended” images in Figure 1c, as observed in data in [12].

**Figure 1:**
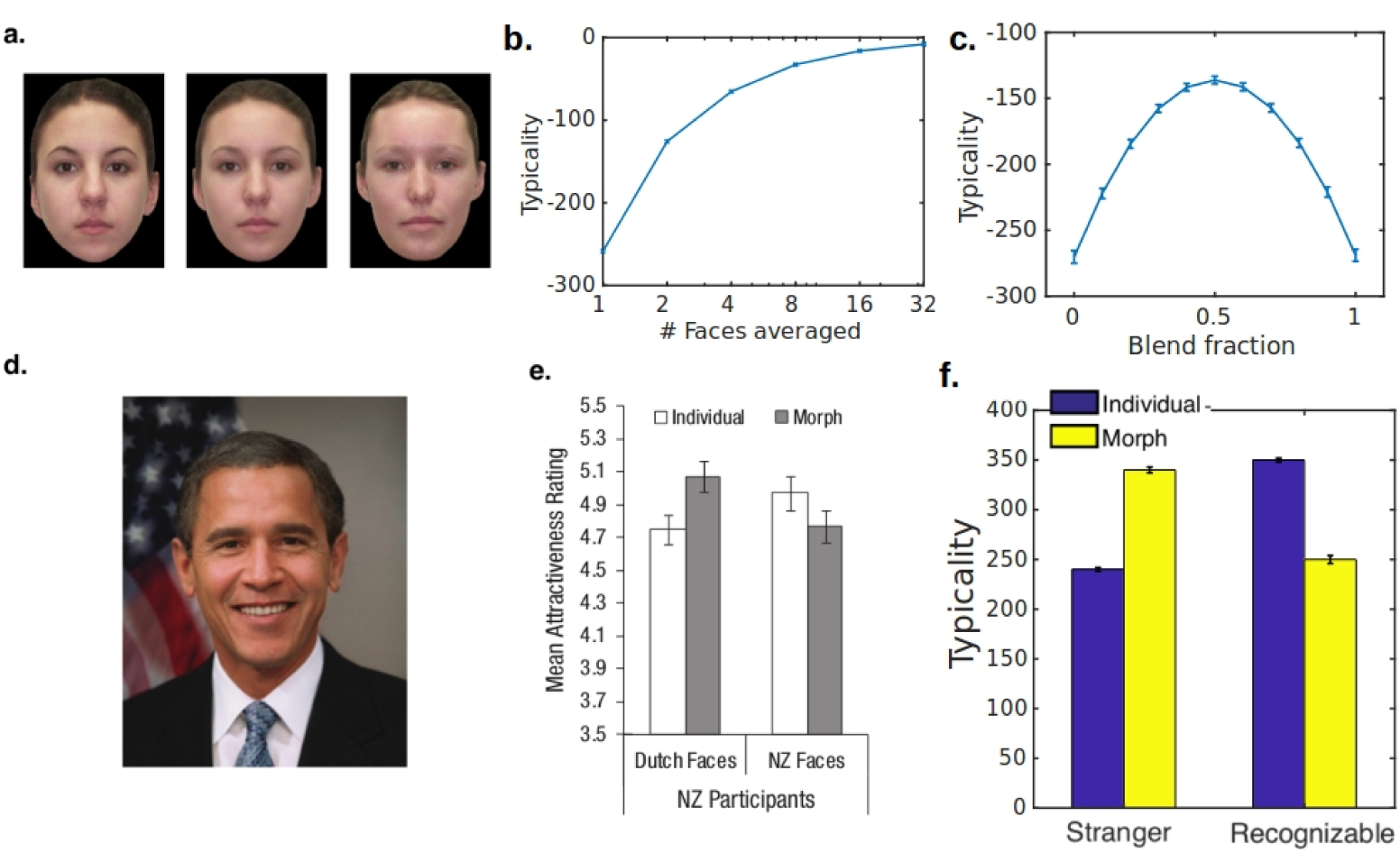
**a.** An example of stimuli (from [12]), with the face in the middle being a 50% blend of the faces on the left and right. **b.** *Model simulation*: a trend of increasing typicality with increasing number of faces used in the composite face. [31] shows a linear trend of composite faces becoming more attractive as more faces were were used in the composite. **c.** *Model simulation*: behaviour of typicality is qualitatively similar to the classical BiA effect: morphs of unknown/stranger faces tend to be rated as more typical than individual constituent faces with the effect becoming more pronounced for more highly blended faces. **d.** An example image (from [11]), depicting a morph of two recognizable faces. **e.** *Data*: (image from [11]) Morphs of recognizable individuals tend to be rated as less attractive than individual, recognizable faces while the morphs of stranger faces tend to be rated as more attractive (BiA). **f.** *Model simulation*: Typicality from model simulation show similar behaviour to mean attractiveness ratings in the data **e.** (A constant offset of 500 was added to make comparison to **e.** easier)

##### UiA: Familiar Faces

In [11], participants from Netherlands and New-Zealand rated morphs of local celebrities (people famous in one country but not the other). Blends of unknown celebrities were rated as more attractive than the constituent images (classic BiA) while blends of local celebrities as less attractive relative to the constituent images: a reversal of BiA. An example image (from [11]) depicting a morph of two recognizable faces can be seen in Figure 1d, while Figure 1e shows BiA and its reversal in data from the study. As discussed in section 3, low statistical typicality of the blend in the distinctive feature subspace results in UiA. Simulations qualitatively capture this effect in Figure 1 **f**.

##### UiA: Race Categorization

In [12], participants rated mixed and single race morphs on attractiveness after performing a race categorization task (Asian or Caucasian). An example of stimuli used in [12] is shown in Figure 2a. Data in Figure 2b shows that mixed race morphs are rated as less attractive relative to single race morphs when a race categorization task preceded the attractiveness judgment. As previously hypothesized in section 3, low statistical typicality of a mixed race blend in the subspace of race informative features induces UiA. For simplicity, we assume this subspace is the one determined by Linear Discriminant Analysis (LDA). Simulations qualitatively capture the behaviour of attractiveness judgments in data in Figure 2**c**.

**Figure 2:**
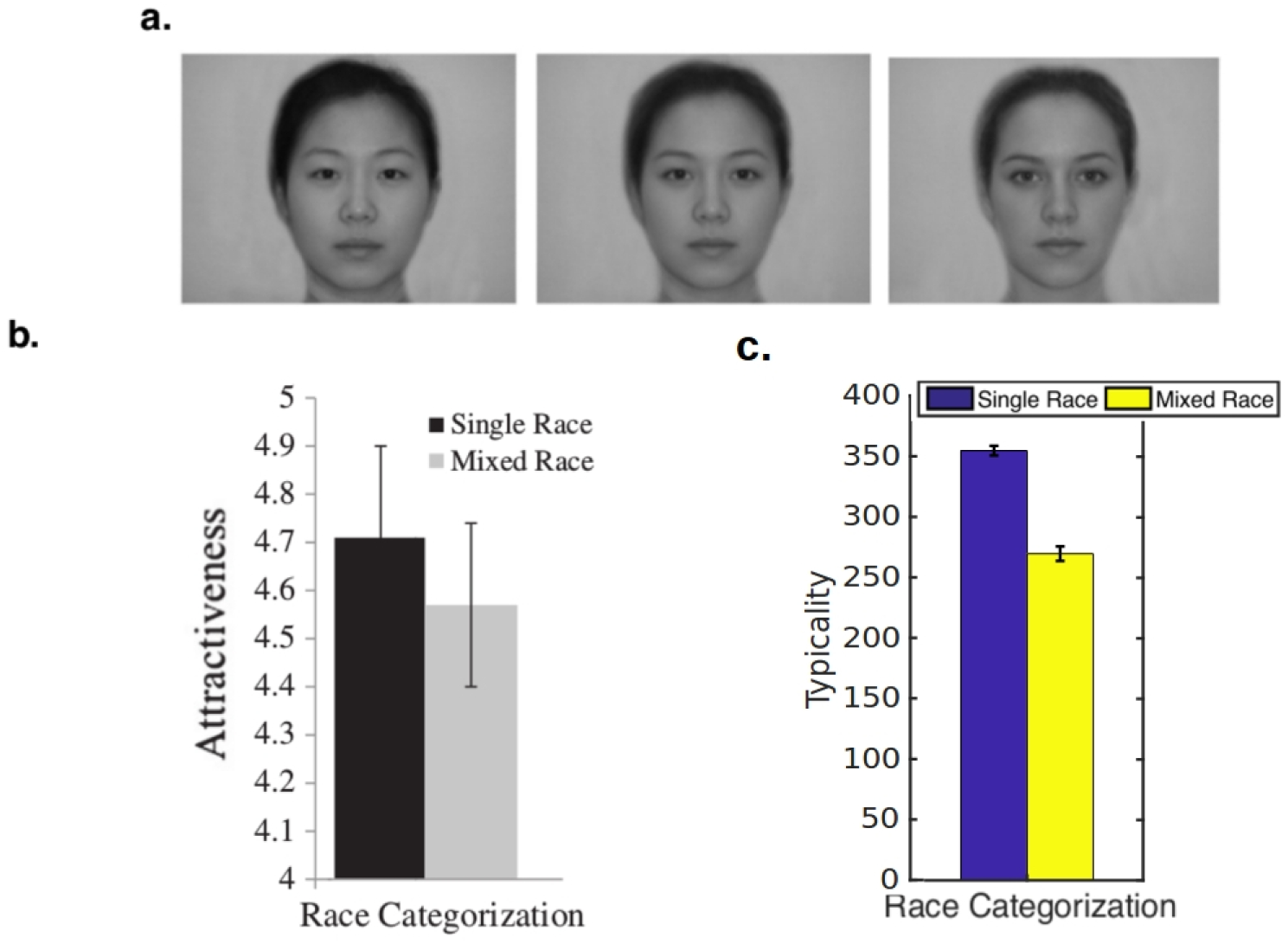
**a.** An example of stimuli used in [12], with the middle face being a 50% blend of the Asian and Caucasian faces shown. **b.** *Data*: (image from [12]) Mean attractiveness ratings for single race and mixed race morphs when a race categorization (Asian or Caucasian) preceded the attractiveness rating. **c.** *Model simulation*: Typicality exhibits qualitatively similar behaviour as human data: mixed race morphs are less typical relative to single race morphs only when a race-categorization precedes the typicality rating (A constant offset of 500 was added to make comparison to **b** easier).

(Simulation parameters: *d* = 100, *d*_race_ = 5, *s* = 10, 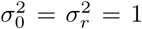 and *μ* = 1. The number of known/recognizable faces |*K*| is set to 50 and the prior probability of a recognizable face 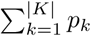 = 0.01.)

### 4.2 Data-Driven Face Representation

Several properties of a vector space representation of faces are desirable: (1) we want a sufficiently complex representation, such that each real face image maps to a unique point in this space; (2) we want it to be a generative model, such that each point generates a realistic face image; (3) we want the dimensionality of the space to be small enough to facilitate useful analyses; (4) an added bonus is if the representation has some neural relevance.

The above desiderata led us to AAM, a well-established machine vision technique that does a good job of reconstructing images, generates realistic synthetic faces and has a latent representation of only a few dozen features [8, 4]. Additionally, it has been shown that AAM features are well encoded by face processing neurons in the monkey brain, and that this neuronal representation is rich enough to discriminate between different individuals [3]. Finally, AAM also has a fairly transparent representation as follows. Each face image defines a *shape vector*, which is just the (*x, y*) coordinates of some consistently defined landmarks across all faces – in our case, we use the free software Face++ (https://www.faceplusplus.com), which labels 83 landmarks (e.g. contour points of the mouth, nose, eyes). Each face image also defines a *texture vector*, which is just the grayscale pixel values of a warped version of the image after aligning the landmark locations to the average landmark locations across the data set. The dataset we use consists of 597 face images with neutral facial expression taken in the laboratory [19]. We learned an AAM that contains a set of basis that describe the joint variations in shape and texture and principal components (PC) representing the variation of the shape and texture of a face from the average face, see Figure 3a for an illustration.

**Figure 3:**
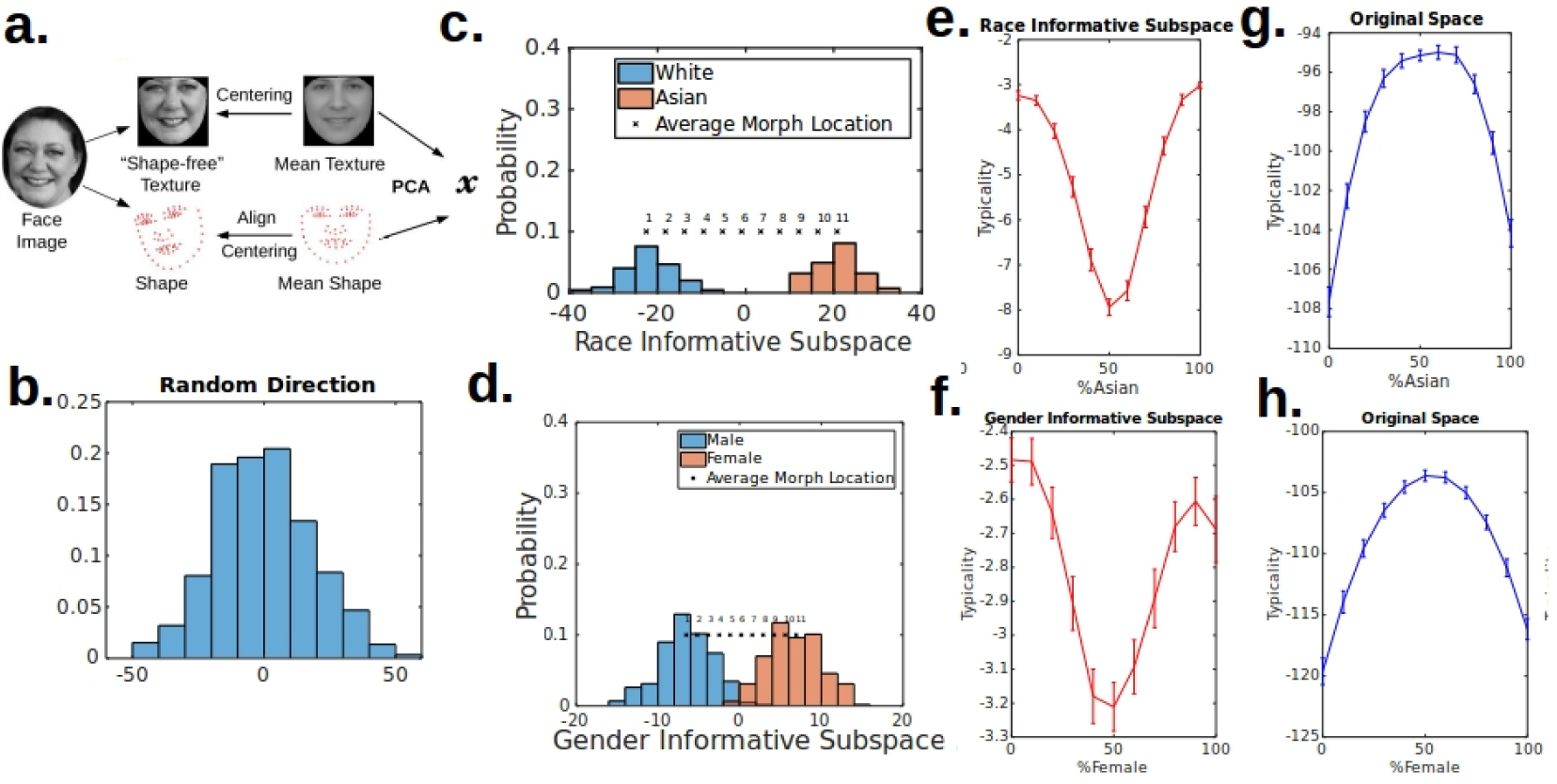
**a.** An illustration of the AAM used for learning the face space and it’s statistical distribution. **b.** The empirical distribution projected in a random direction/1-d subspace is normally distributed. **c, d** The empirical distribution of faces projected in race or gender informative subspace show that the distribution is a mixture of normals. Also shown are the average locations of the continuum of race and gender blends used in the simulation. **e, f** Statistical typicality in category informative subspace induces UiA. **e, f** Statistical typicality in original/full space induces BiA, when there is no categorization task.

#### Simulations

First, we validate several assumptions we made in our abstract model. In Figures 3b;c;d, it can be seen that the distribution of faces learned from data is normally distributed in random directions but can be seen to be a mixture of normal distributions in race or gender informative subspaces (here, found using LDA). In Figures 3e;f;g;h, it can be seen that statistical typicality in categorization informative subspace induces UiA; the same stimuli induce BiA in original/full space or random subspace (not shown), which would be the case in the absence of the categorization task. These results also highlight the importance of attentional modulation of face space.

## 5 Experimental Validation

In [28], participants rate the attractiveness of blends on a male-female continuum following a gender categorization, see Figure 4a. In the control condition, participants only rated attractiveness of the same images without categorization. We projected the stimuli into our face space learned using the AAM, see Figure 4b and computed statistical typicality in the original/full space as well as the task informative subspace, see Figure 4c;d. We found that typicality in the task informative subspace is significantly correlated (*p* < 0.05) with the difference in attractiveness ratings between the experiment and control condition, consistent with our model.

**Figure 4:**
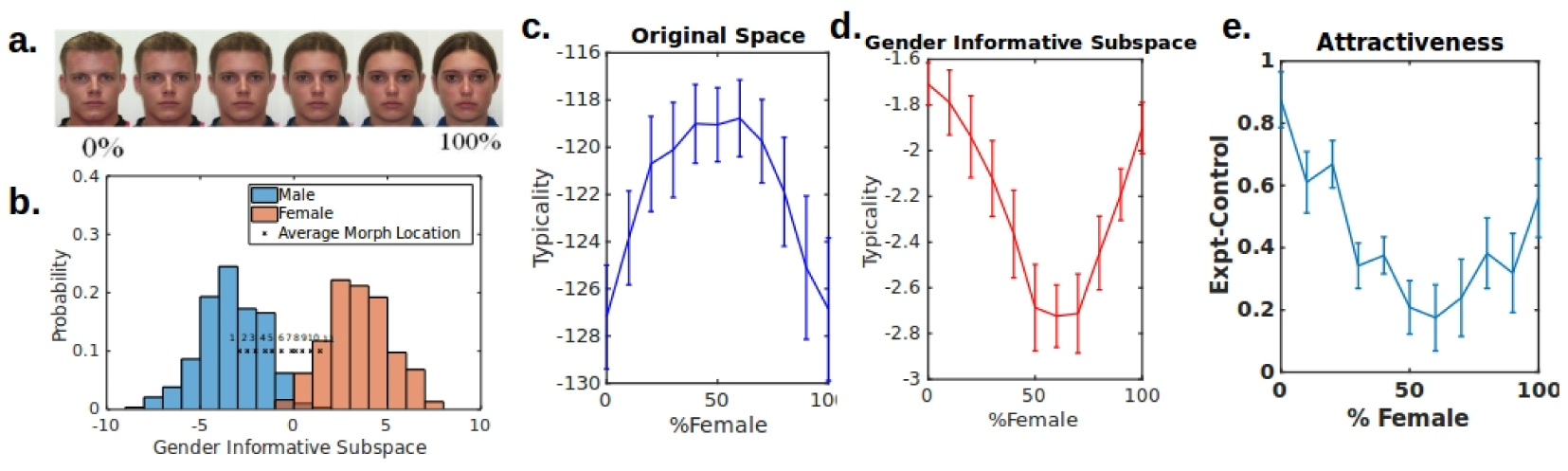
**a.** An example of stimuli used in [28]: a continuum of male-female gender blends. **b** Analogous to Fig. 3c;d but projecting the actual contiuum of gender blends used as stimuli. **c, d** Analogous to Fig. 3 e;f;g;h using experiment stimuli **e** Difference in attractiveness ratings between experiment and control condition shows similar trend as in **d** as expected.

## 6 Disentangling the Disfluency and Typicality accounts

In the simulations and experiments considered so far, our statistical typicality account and the disfluency account make similar qualitative predictions, though the disfluency account is unable to make quantitative predictions. To disambiguate these accounts of UiA, we propose an experiment in which these two accounts make different predictions. In Figure 5a, we plot the empirical distribution of ages in a publicly available dataset [1]. Participants can be trained to learn this distribution [6] (though not via categorization to avoid categorical percept [18]). The proposed task is to rate attractiveness of faces after an age categorization task: older or younger than 37.5 y? According to the statistical typicality account, attractiveness ratings would look like Figure 5b. While the disfluency account does not make any quantitative predictions, we expect that faces close to 37.5 in age will be difficult to classify and take longer and are expected to induce UiA, while the typicality account is expected to induce BiA.

**Figure 5:**
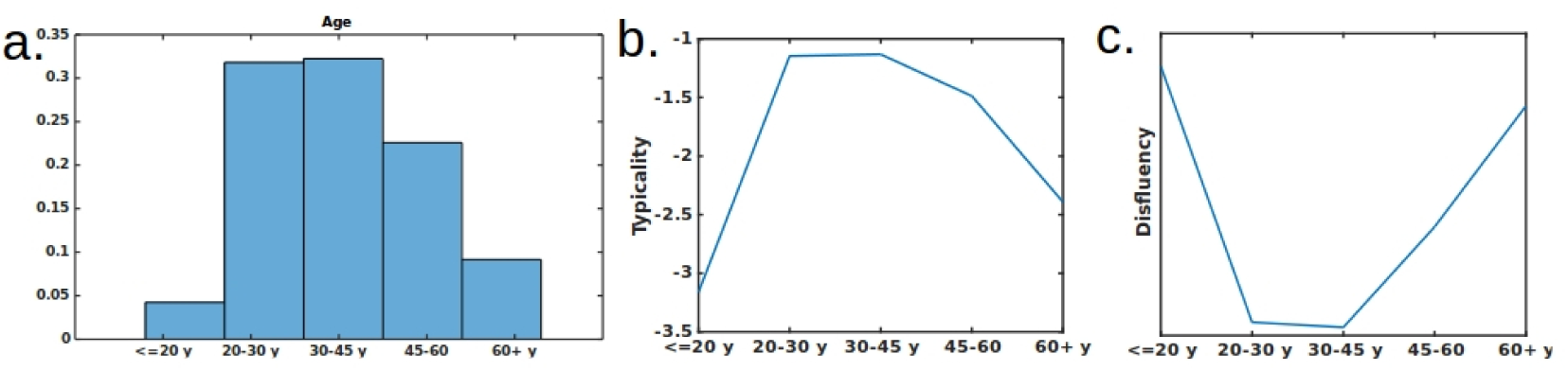
**a.** Empirical distribution of ages in dataset [1]. **b** Predictions of typicality based model. **c** Expected predictions based on a disfluency account.

## 7 Discussion

Most existing models of human preferences assume these preferences to be fixed and do not model contextual dependence. In this paper, we propose a statistical model of “liking” that assumes it is positively influenced by perceived typicality that is context sensitive. We assume that humans naturally project high-dimensional data, such as faces, to a low-dimensional subspace representation, either via bottom-up saliency (distinctiveness with respective to the assumed generative distribution) or via top-down goal-directed specification (informativeness with respect to a particular discrimination problem, such as race or gender). The bottom-up saliency component explains the celebrity-morph data, while the top-down component explains the influence of racial/gender discrimination on attractiveness perception.

In addition to providing a statistically grounded explanation of contextual dependence of human attractiveness judgment, our work also provides some general insight as to how high-dimensional data can be analyzed and stored efficiently in a low-dimensional representation, as long as the system is equipped to dynamically shift its subspace projection according to task demands. This approach may well find applications in modeling other types of human preferences in high-dimensional data space, and explain context sensitivity in those domains. The general idea of “tagging” high-dimensional data by their distinctive features seems like a good way to store and analyze complex data. Future work is needed to investigate the theoretical and numerical consequences of using this approximate representation.

This work also sheds light on one possible statistical role played by attention. It is one way to dynamically construct subspaces that emphasizes feature dimensions that are most relevant or salient for performing the task at hand. There is a broad and confusing literature of attention in both psychology and neuroscience. A productive direction of future research would be to relate our hypothesized role of attention here to that large literature.

## 8 Acknowledgments

We thank Piotr Winkielman and Jamin Halberstadt for sharing the gender categorization data and Samer Sabri for writing input.

